# Benchmarking Transposable Element Annotation Methods for Creation of a Streamlined, Comprehensive Pipeline

**DOI:** 10.1101/657890

**Authors:** Shujun Ou, Weija Su, Yi Liao, Kapeel Chougule, Doreen Ware, Thomas Peterson, Ning Jiang, Candice N. Hirsch, Matthew B. Hufford

## Abstract

Sequencing technology and assembly algorithms have matured to the point that high-quality *de novo* assembly is possible for large, repetitive genomes. Current assemblies traverse transposable elements (TEs) and allow for annotation of TEs. There are numerous methods for each class of elements with unknown relative performance metrics. We benchmarked existing programs based on a curated library of rice TEs. Using the most robust programs, we created a comprehensive pipeline called Extensive *de-novo* TE Annotator (EDTA) that produces a condensed TE library for annotations of structurally intact and fragmented elements. EDTA is open-source and freely available: https://github.com/oushujun/EDTA.

## Background

Transposable elements (TEs) are repetitive, mobile sequences found in most eukaryotic genomes analyzed to date. Originally discovered by Barbara McClintock in maize (*Zea mays*) [1], TEs are now known to comprise the majority of genetic material in many eukaryotes. For example, TEs make up nearly half of the human (*Homo sapiens*) genome [2] and approximately 85% of the genomes of wheat (*Triticum aestivum*) and maize [3, 4]. The functional and evolutionary significance of TEs has also become increasingly clear. *Stowaway* and *PIF*/*Harbinger* transposons in rice (*Oryza sativa*), for instance, are associated with subspecies-specific hotspots of recombination [5], and specific TE insertions have been associated with plant architecture [6] and flowering time [7] in maize, generating phenotypic variation important during domestication and temperate adaptation.

Despite their prevalence and significance, TEs have remained poorly annotated and studied in all but a few model systems. Transposable elements create a particularly challenging genome assembly problem due to both their high copy number and the complex nesting structures produced by new TE insertions into existing TE sequences. While the low-copy, genic fraction of genomes has assembled well, even with short-read sequencing technology, assemblies of TEs and other repeats have remained incomplete and highly fragmented until quite recently.

Long-read sequencing (*e.g.*, PacBio and Oxford Nanopore) and assembly scaffolding (*e.g.*, Hi-C and BioNano) techniques have progressed rapidly within the last few years. These innovations have been critical for high-quality assembly of the repetitive fraction of genomes. In fact, Ou *et al.* [8] demonstrated that the assembly contiguity of repetitive sequences in recent long-read assemblies is even better than traditional BAC-based reference genomes. With these developments, inexpensive and high-quality assembly of an entire genome is now possible. Knowing where features (*i.e.*, genes, TEs, etc.) exist in a genome assembly is important information for using these assemblies for biological findings. However, unlike the relatively straightforward and comprehensive pipelines established for gene annotation [9–11], current methods for TE annotation can be piecemeal, inaccurate, and are highly specific to classes of transposable elements.

Transposable elements fall into two major classes. Class I elements, also known as retrotransposons, use an RNA intermediate in their “copy and paste” mechanism of transposition [12]. Class I elements can be further divided into long terminal repeat (LTR) retrotransposons, as well as those that lack LTRs (non-LTRs), which include long interspersed nuclear elements (LINEs), and short interspersed nuclear elements (SINEs). Structural features of these elements can facilitate automated *de novo* annotation in a genome assembly. For example, LTR elements have a 5-bp target site duplication (TSD), while non-LTRs have either variable length TSDs or lack TSDs entirely, being instead associated with deletion of flanking sequences upon insertion [13]. There are also standard terminal sequences associated with LTR elements (*i.e.*, 5’-TG…C/G/TA-3’ for LTR-*Copia* and 5’-TG…CA-3’ for LTR-*Gypsy* elements), and non-LTRs often have a terminal poly-A tail at the 3’ end of the element (see [14] for a complete description of structural features of each superfamily).

The second major class of TEs, Class II elements, also known as DNA transposons, use a DNA intermediate in their “cut and paste” mechanism of transposition [15]. As with Class I elements, DNA transposons have superfamily-specific structural features that can be used to facilitate an automated identification process [16]. For example, *hAT* elements typically have an 8-bp TSD, 12-28 bp terminal inverted repeat sequence (TIRs), and contain 5’-C/TA…TA/G-3’ terminal sequences. Each Class II superfamily has different structural features that need to be considered when TE annotation programs are being developed and deployed [16, 17]. *Helitrons* are a unique subclass of Class II elements that replicate through a rolling-circle mechanism and, as such, do not generate a TSD sequence and do not have TIRs, but do have a signature 5’-TC…CTRR-3’ terminal repeat sequence and frequently a short GC-rich stem-loop structure near the 3’ end of the element [16,18,19].

High-quality TE annotations have been generated for several model species through extensive community efforts and manual curation (*e.g.*, human [2], *Drosophila melanogaster* [20], *Arabidopsis thaliana* [21], rice [22, 23], maize [4], etc.). However, with numerous reference genome assemblies being generated both within and across species, manual curation is no longer feasible, and automated annotation of TEs is required. Dozens of programs have been developed for this purpose and these generally fall into one of three categories [24, 25]. First, general repeat finders identify high copy number sequences in a genome [26–28]. These programs can have high sensitivity for identifying repetitive sequences, but have limited ability to classify them into specific TE superfamilies and can misidentify non-TE features (*e.g*., high copy-number genes). Second, the sequence homology approach [29–32] is quick and takes advantage of prior knowledge (*i.e.*, databases), but is limited by the depth and accuracy of this knowledge and variability across TE sequences. The final approach takes advantage of the structural makeup of classes and superfamilies of TEs for *de novo* structural annotation [24, 25]. This approach is advantageous in that it is codable and does not rely on repeat databases, therefore being ideal for new species. However, the approach is limited by the knowledge of the sequence structure of TEs, and is often characterized by a high false discovery rate.

While numerous and, in some cases, redundant TE identification methods exist, their performance has not been comprehensively benchmarked, despite recognition that this would be an important exercise [33]. Here, we have gathered a broad set of existing TE annotation software and, using several metrics, have compared each program’s performance to a highly curated TE reference library in rice. Based on our benchmarking results, we propose an optimal pipeline for the generation of *de novo* TE libraries that can then be used for genome annotation.

## Results

In eukaryotic genomes, transposable elements (TEs) are present in both structurally intact and fragmented sequences. In addition, extensive sequence variation among species can make annotation of these sequences challenging. Development of a species-specific TE library is an essential step in the annotation process, which begins with structural identification of major TE classes followed by manual curation. Representative sequences, or exemplars, representing their respective TE families, are then used to detect fragmented and mutated TE sequences that are not recognizable using structural features. Importantly, if there are errors in the annotation library, these will be propagated during the whole-genome annotation process.

A number of computer programs are available for structural identification of TEs and development of *de novo* annotation libraries. We have benchmarked commonly applied programs for metrics including sensitivity, specificity, accuracy, and precision (Figure 1) and have implemented the optimal set of programs in a comprehensive pipeline called the Extensive *de-novo* TE Annotator (EDTA). To evaluate each program, we used a high-quality, manually curated library developed for the model species *Oryza sativa* (rice), which has a long history of TE discovery and annotation [23,34–42]. The reference library contains abundant class I and class II elements, making it optimal for testing currently available annotation programs.

**Figure 1.**
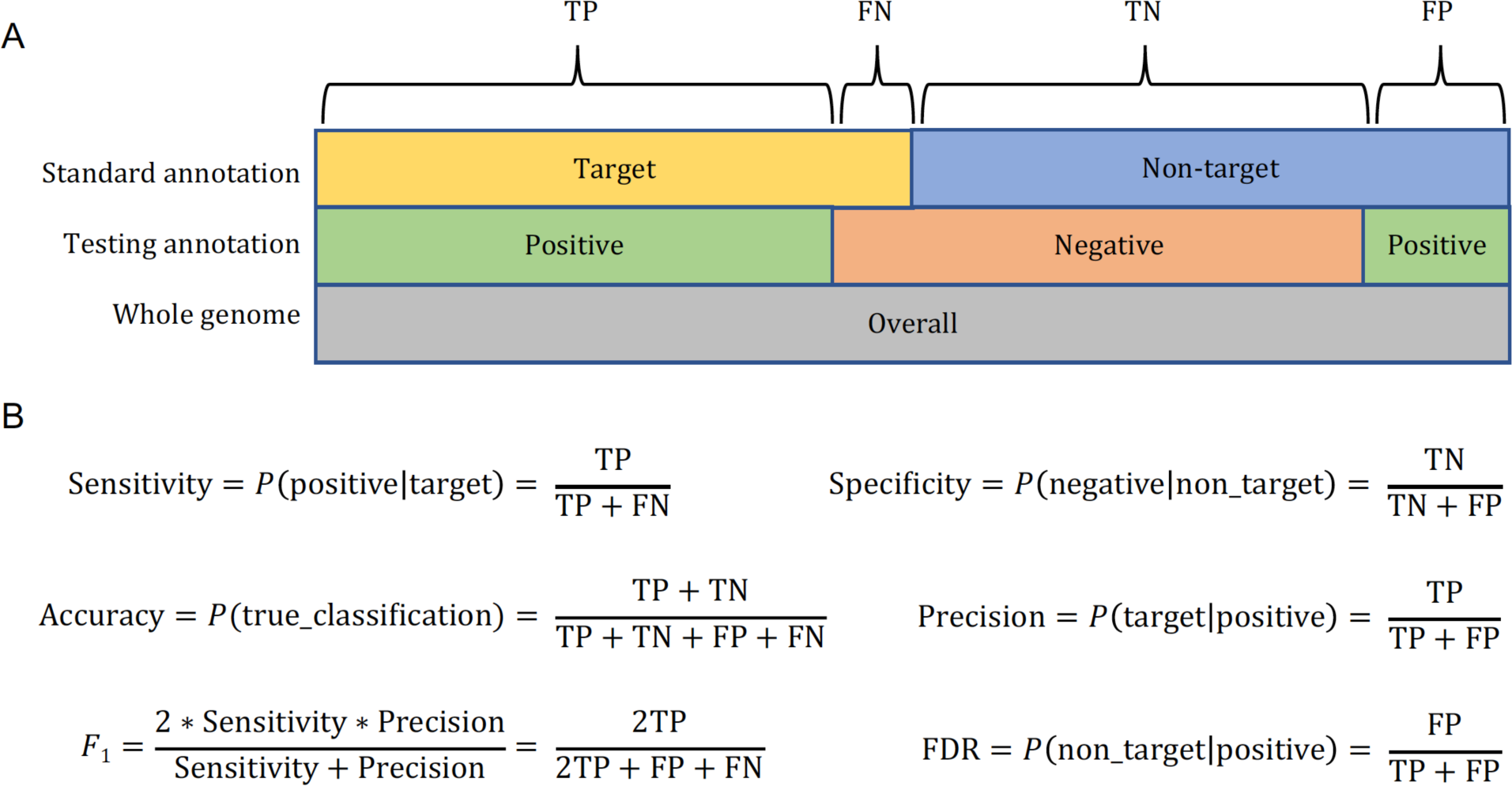
Schematic representation of benchmarking metrics. (A) Definition of TP, true positive; FP, false positive; FN, false negative; TN, true negative. (B) Definition of sensitivity, specificity, accuracy, precision, *F*_1_ measure, and FDR. Each metric is calculated based on genomic sequence length in bp.

### Setting up a reference annotation for benchmarking

The reference annotation library for rice was created through substantial manual curation of repeat families obtained from an all-versus-all BLAST search of the rice genome (details in Methods). This library was then used to annotate the rice genome for both structurally intact and fragmented TE sequences, which comprised 23.98% and 22.66% of the rice genome, respectively (46.64% in total; Table 1). Since half of all TEs in the rice genome are fragmented, structural annotation alone would miss a substantial portion of TE sequences. A homology-based approach that uses a TE library is necessary to obtain a complete annotation.

**Table 1.** TE content in the rice (*Oryza sativa* ssp. *japonica* cv. ‘Nipponbare’ v. MSU7) genome.

|  | Class | Std6.9.5* | Complete** | Fragmented** | Total** |
| --- | --- | --- | --- | --- | --- |
| LTR | Class I | 88.1 Mb | 14.44% | 9.11% | 23.54% |
| non-LTR | Class I | 7.6 Mb | 0.51% | 1.52% | 2.03% |
| TIR | Class II | 65.5 Mb | 7.93% | 9.56% | 17.49% |
| *Helitron* | Class II | 13.4 Mb | 1.10% | 2.47% | 3.57% |
| Total | - | 174.6 Mb | 23.98% | 22.66% | 46.64% |

\* Annotation based on the curated library (v6.9.5).
\*\* Percent of genome estimated based on a genome size of 374.3 Mb.

TEs in this annotation library are broken down into a number of non-overlapping categories, including LTR (referring to LTR retrotransposons), non-LTR (including SINEs and LINEs), TIR (referring to DNA transposons with TIRs, including MITEs), *Helitron*, and non-TE repeat sequence. LTR retrotransposons contribute the largest component, 23.54% of the total genomic DNA (Table 1). Non-LTR retrotransposons including SINEs and LINEs contribute the smallest proportion of total sequence (7.6 Mb or ∼2% of the genome; Table 1). DNA transposons contribute ∼21% (17.49% TIR elements and 3.57% *Helitrons*; Table 1).

To test various programs, the genome was partitioned into target and non-target sequences (Figure 1A). For example, when testing the performance of an LTR annotation program, LTR sequences matching our curated library were labeled “target” and all other sequences were labeled “non-target”. Each program’s annotation was then compared to that from our curated library, with sequences included in our target subset counted as true positives (TP), sequences in our non-target subset categorized as false positives (FP), missed targets counted as false negatives (FN), and the remainder of the genome (not TP, FP, nor FN) labeled as true negative (TN) (Figure 1A).

We then used six metrics (sensitivity, specificity, accuracy, precision, FDR, and *F*_1_) to characterize the annotation performance of the test library created by various programs (Figure 1B). These metrics were calculated based on the total number of genomic DNA bases, because misannotations occurring in the test library will be amplified in the whole-genome annotation process. *Sensitivity* denotes how well the test library can correctly annotate target TE sequences. *Specificity* describes how well the test library can correctly exclude non-target sequences. *Accuracy* denotes the true rate in discriminating target and non-target sequences. *Precision* is the true discovery rate, while *FDR* is the false discovery rate. Finally, the *F*_1_ measure is the harmonic mean of precision and sensitivity; *F*_1_ is similar to accuracy, but is useful because it does not require an estimate of TN, which can be difficult to quantify. While we can estimate TNs with the use of the curated annotation, we still include the *F*_1_ measure in our study to allow for comparison to previous work.

We exhaustively searched the literature for open-source programs and databases that have been developed for general repeat annotations as well as structural annotation programs for LTR elements, SINEs, LINEs, TIR elements, and *Helitrons*. We applied educated parameters based on knowledge of transposon structures to run these programs (see Methods and Additional File 1). We also applied filters on initial program predictions to remove low-quality candidates and potentially false predictions such as short sequences and tandem-repeat-containing sequences (Additional File 1). For each program, a non-redundant test library was created from filtered TE candidates, which was then used to annotate the rice genome. The testing annotation from each program for each category of TEs was compared with the annotations from the curated library for calculation of benchmarking metrics.

### Comparison of General Repeat Annotators

We benchmarked five general repeat annotators, including RECON [43], RepeatScout [26], RepeatModeler [28], Red [27], and Generic Repeat Finder (GRF) (https://github.com/bioinfolabmu/GenericRepeatFinder), as well as a repeat database Repbase [30], which is widely used as the default library in RepeatMasker [29]. For these TE annotation approaches, only RepeatModeler and Repbase provide classification of annotated TEs. Among these methods, we found that Repbase employing the rice TE database had very high performance in both TE identification and classification (Figure 2), which is a product of continuous improvement and curation of rice TEs by the community. However, if we exclude rice related TEs in Repbase and treat rice as a newly sequenced species (Repbase_norice in Figure 2), the annotation (Figure 2A) and classification (Figure 2B) sensitivity both drop from ∼94% to ∼29%, despite extremely high specificity (∼99%) and low FDR (∼5%) (Table S1A). This result was consistent for each of the TE classes (Figure 3A – LTR elements; Figure 4A – non-LTR elements; Figure 5A – TIR elements; Figure 6A – *Helitron*), though the drop in sensitivity was substantially greater for *Helitrons* (to less than 5%) than for other elements. For TE classifications, RepeatModeler performed similarly to Repbase without rice sequences (Figure 2B), and both can, therefore, be used as high-quality supplements to other specialized TE annotators. GRF is the most recently developed general repeat finder, but had the lowest sensitivity (75%), which is likely due to its inability to introduce gaps during the multiple sequence alignment process (Table S1A).

**Figure 2.**
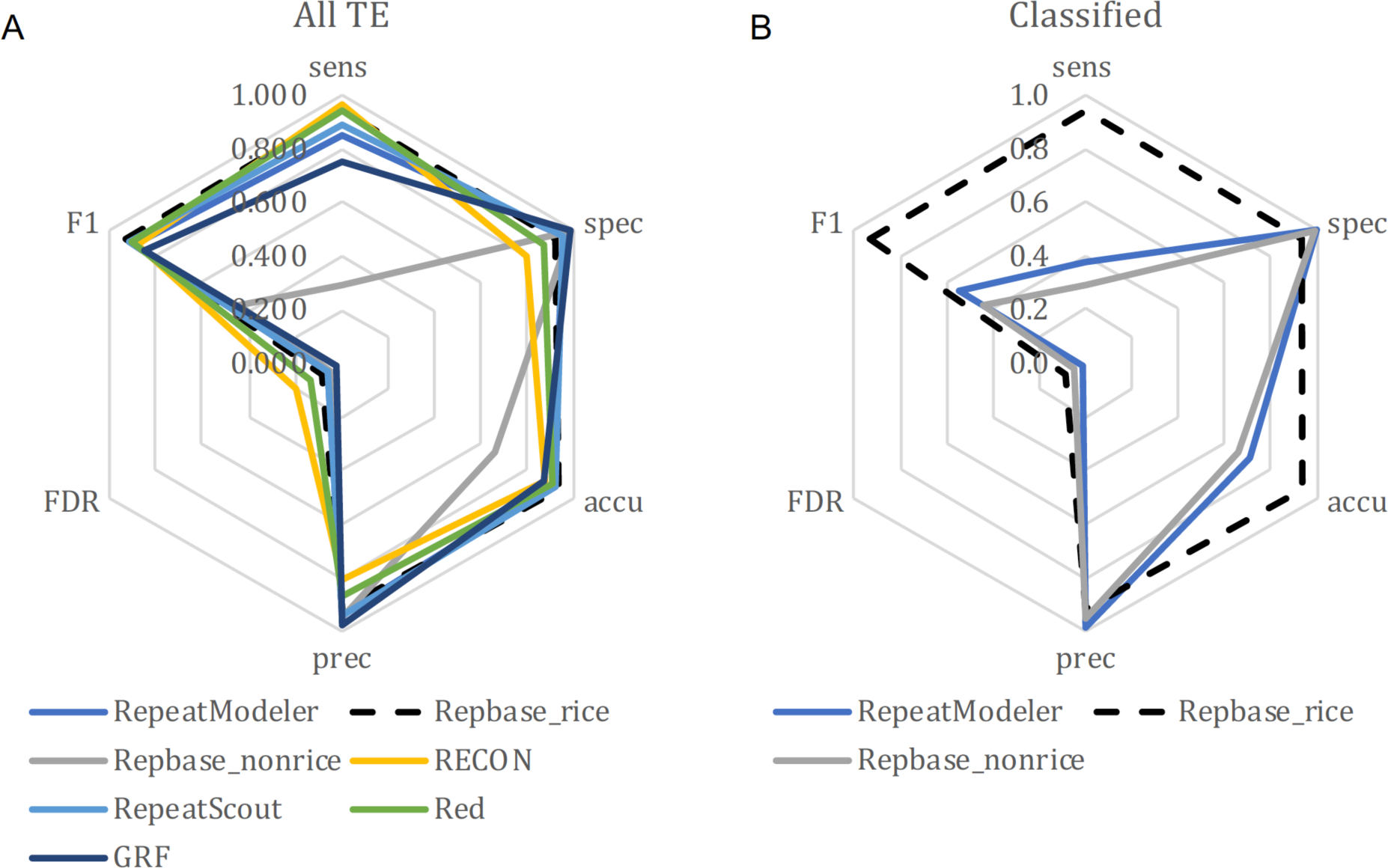
Annotation performance of general repeat annotators compared to the rice curated annotation. (A) Annotation and (B) Classification performance of various methods. Sens, sensitivity; Spec, specificity; Accu, accuracy; Prec, precision; FDR, false discovery rate; F1, *F*_1_ measure.

**Figure 3.**
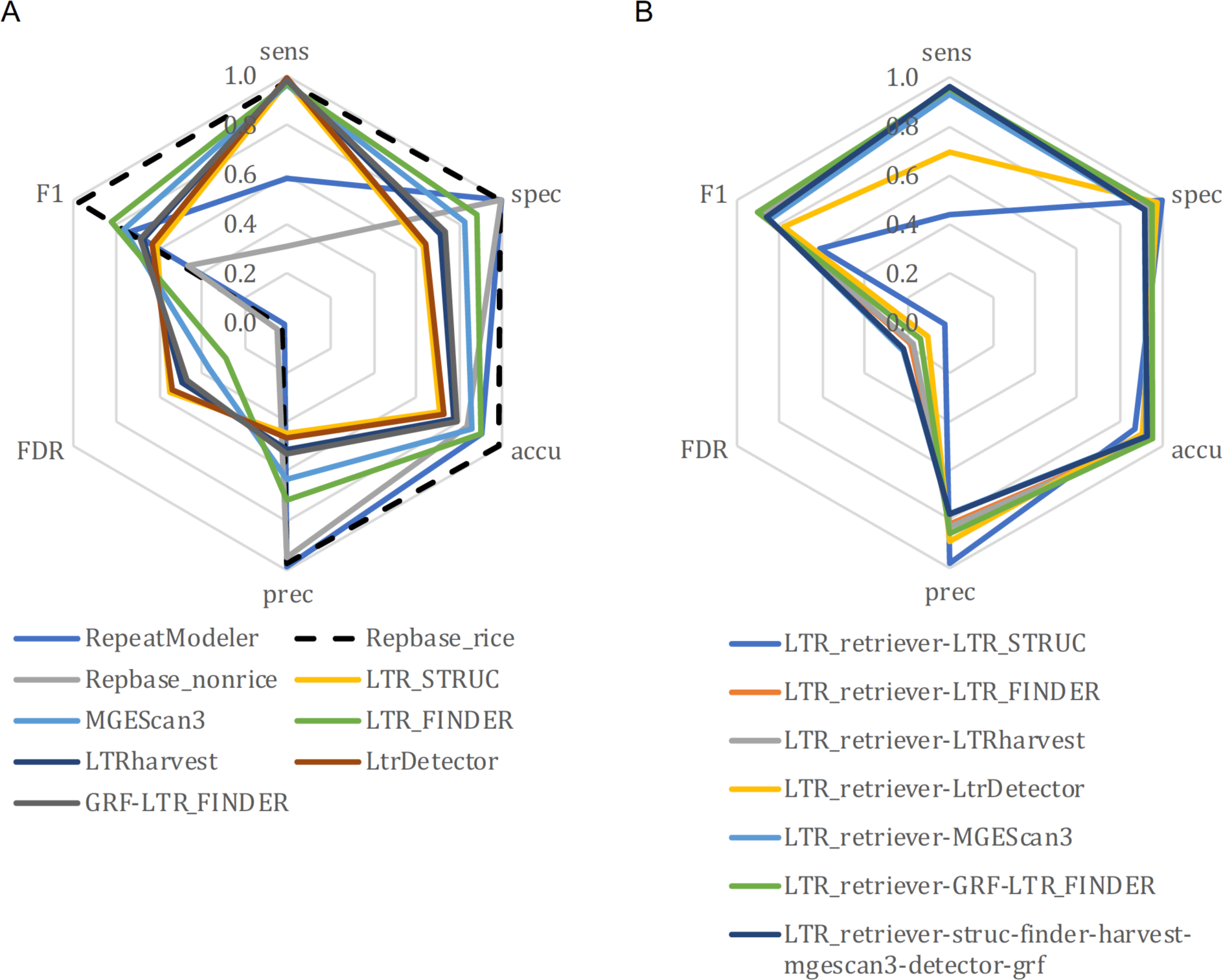
Annotation performance of LTR-related programs as compared to the rice curated annotation. (A) Various methods to identify LTR retrotransposons. GRF-LTR_FINDER combines the terminal direct repeat search engine in GRF and the filtering engine in a modified version of LTR_FINDER for detection of LTR retrotransposons. The LTR_FINDER result was generated by the parallel version. (B) LTR_retriever specific results, which were generated using LTR_retriever to process results from other programs specified in each of the names in the figure. Sens, sensitivity; Spec, specificity; Accu, accuracy; Prec, precision; FDR, false discovery rate; F1, *F*_1_ measure.

**Figure 4.**
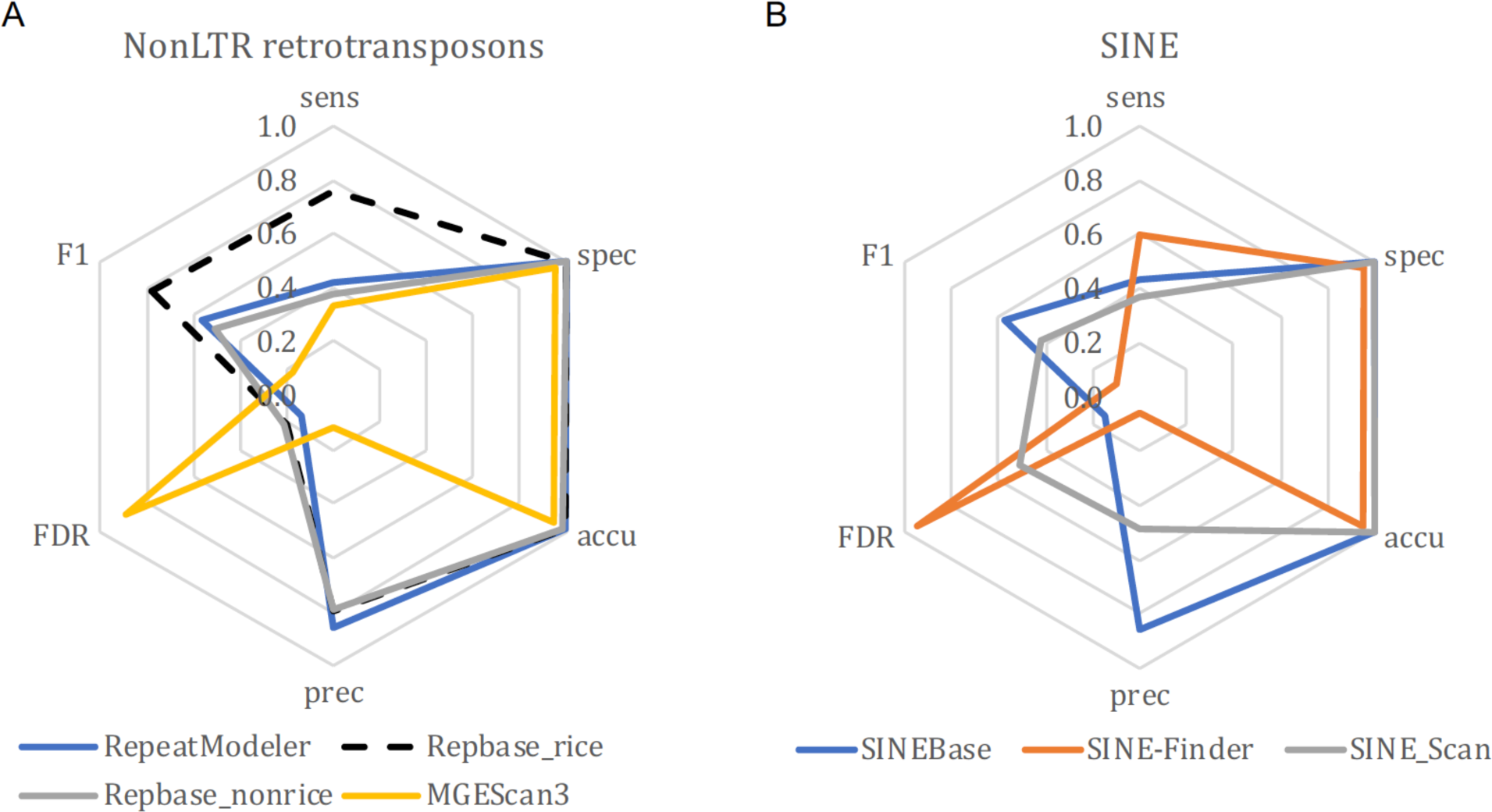
Annotation performance of methods to identify non-LTR retrotransposons as compared to the rice curated annotation. (A) Non-LTR retrotransposon annotation methods. (B) Short interspersed nuclear element (SINE) annotation methods. Sens, sensitivity; Spec, specificity; Accu, accuracy; Prec, precision; FDR, false discovery rate; F1, *F*_1_ measure.

**Figure 5.**
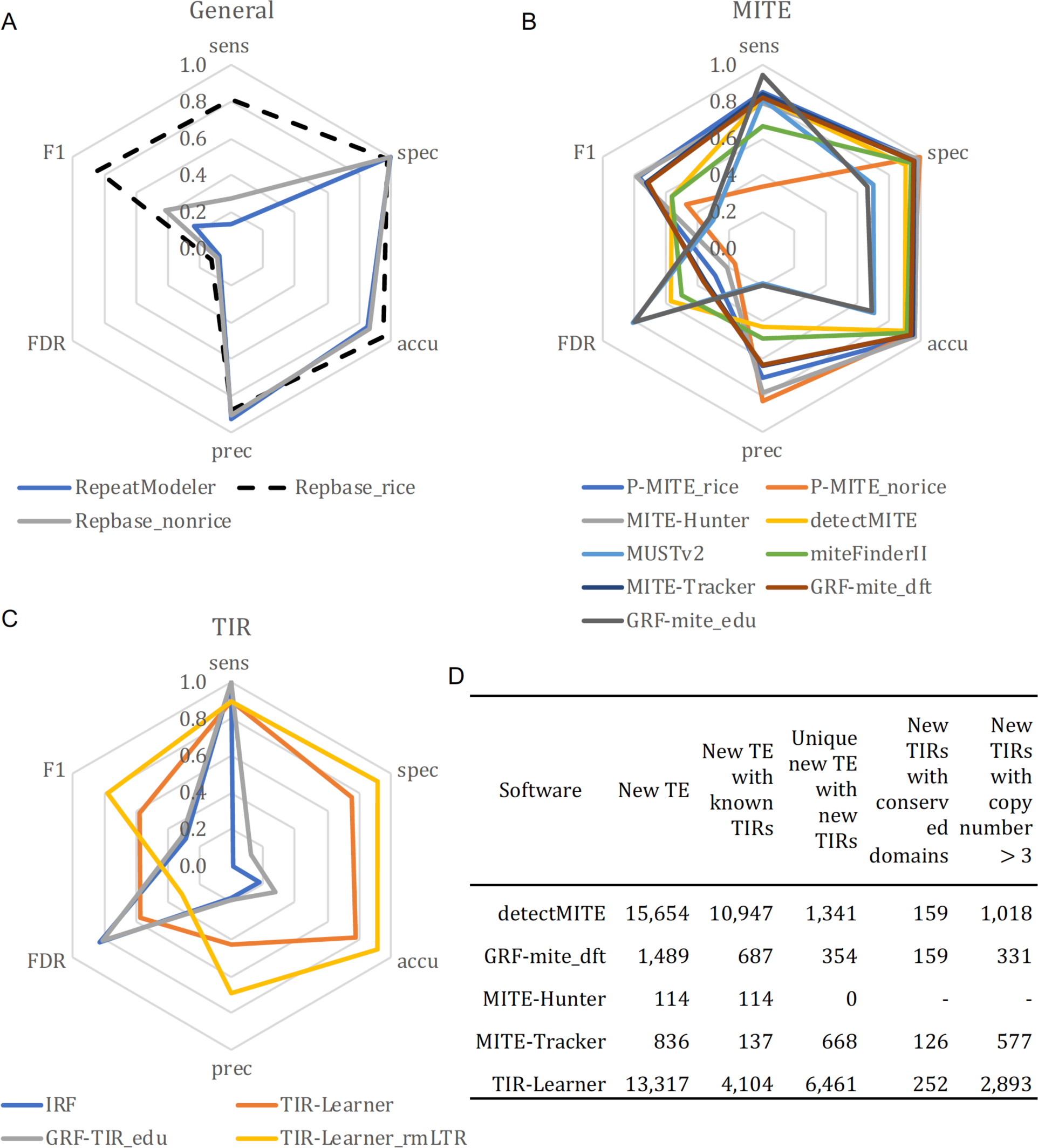
Annotation performance of methods to identify terminal inverted repeats compared to the rice curated annotation. (A) General methods and (C) structure-based methods to identify TIR elements. The TIR-Learner_rmLTR library had LTR-related sequences removed using the curated library. (B) Structure-based methods and specialized database to identify miniature inverted transposable elements (MITEs). Sens, sensitivity; Spec, specificity; Accu, accuracy; Prec, precision; FDR, false discovery rate; F1, *F*_1_ measure. (D) Verification of new TIR candidates identified by TIR-Learner and MITE programs. Conserved protein domains from known TIR elements were used to annotate new TIR candidates.

**Figure 6.**
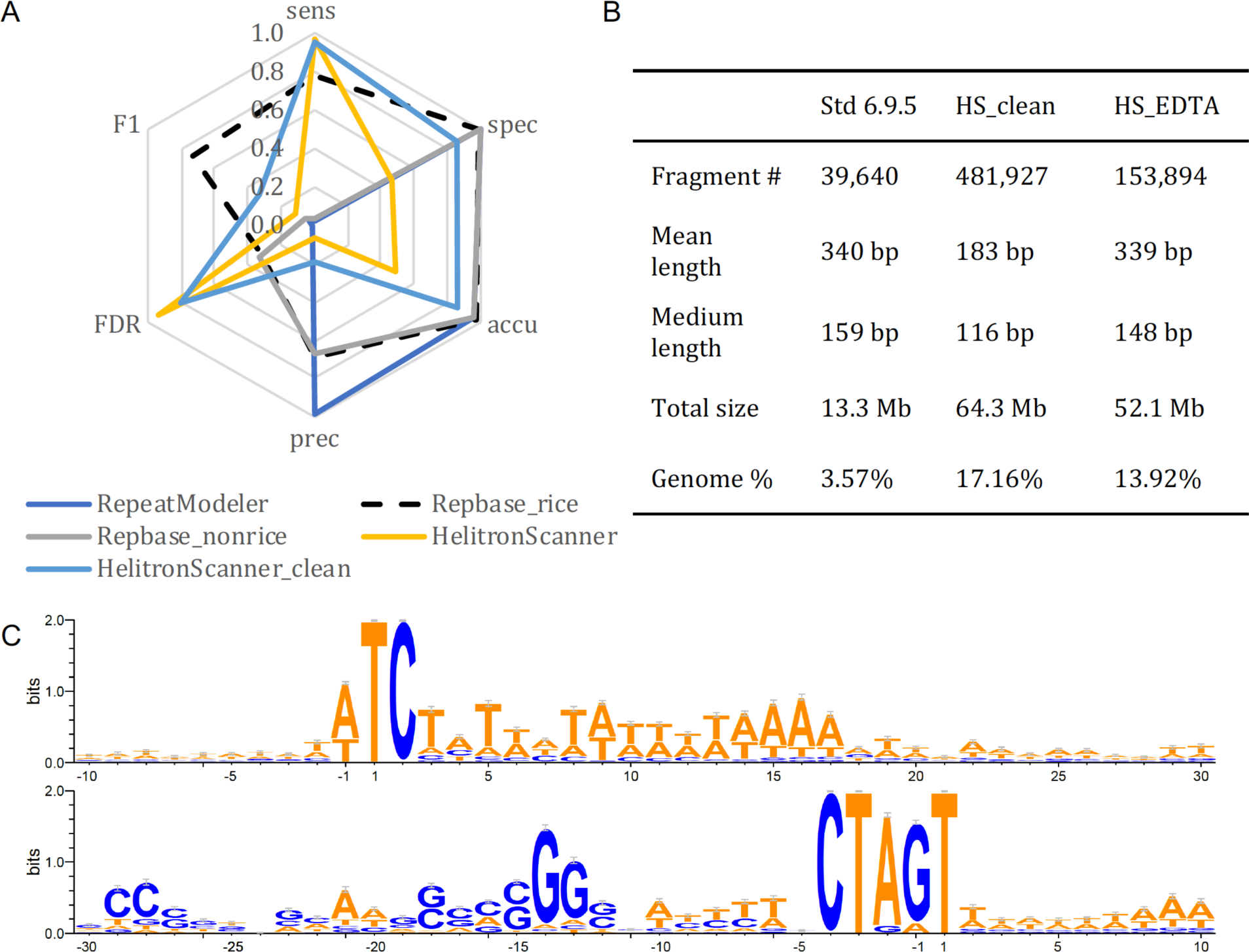
Annotation and characterization of *Helitrons* in rice. (A) Annotation performance of *Helitron*-related methods as compared to the rice curated annotation. The HelitronScanner_clean result had non-*Helitron* TE sequences removed using the curated library. Sens, sensitivity; Spec, specificity; Accu, accuracy; Prec, precision; FDR, false discovery rate; F1, *F*_1_ measure. (B) Comparison of whole-genome annotation results using the curated library (v6.9.5), the HelitronScanner clean library (HS_clean), and the EDTA filtered HelitronScanner library (HS_EDTA). (C) Sequence logos of terminal and flanking sequences of clean *Helitron* candidates. Upper panel, 5’ terminal of *Helitron*-candidates with the starting position labeled as 1; Lower panel, 3’ terminal of *Helitron*-candidates with the last position labeled as −1.

Overall, the general repeat finders we tested have consistently high performance in identifying repetitive sequences in the rice genome, with the exception of Repbase without rice sequences (Figure 2A). What really differentiates these programs is their ease in processing raw results. All are open source and easy to install except Repbase, which requires an institutional subscription for access (Table S2). Red runs on a single CPU and took the shortest time for execution (∼33 min); however, Red produces the largest raw result file, which is highly redundant (35 Mb after clustering; Table S2). RepeatModeler and RepeatScout produced very compact outputs (< 4 Mb). The RepeatScout program runs more efficiently but provides no classification of repeat sequences (Table S2). The RECON and RepeatScout packages are not actively maintained, but have been incorporated into the RepeatModeler package. In summary, RepeatModeler has the highest performance among the general repeat annotators based on our evaluation metrics (Figure 2) and is open source, able to produce a compact output, and able to classify TE families to a certain degree. Still, further classification or use of more specialized software based on the specific structures of each superfamily of TEs is necessary to achieve more accurate annotations.

### Comparison of LTR Annotators

Due to their abundance in eukaryotic genomes, LTR retrotransposons have received the most attention in TE annotation software development. In addition to the two general repeat identification methods with classification (RepeatModeler and Repbase), we found seven structure-based methods that are specifically designed for *de novo* LTR identification. Chronologically in order of development, they are LTR_STRUC [44], LTR_FINDER [45], LTRharvest [46], MGEScan3 [47], LTR_retriever [39], LtrDetector (https://github.com/TulsaBioinformaticsToolsmith/LtrDetector), and GRF (https://github.com/bioinfolabmu/GenericRepeatFinder). In a previous study [39], we developed LTR_retriever and compared its performance to LTR_STRUC, LTR_FINDER, LTRharvest, and MGEScan_LTR [48]. Here, we update the comparison with the recently developed MGEScan3, LtrDetector, and GRF. Meanwhile, the LTR_retriever package has been updated from v1.6 to v2.6 since its initial publication.

The six structural-based methods that we tested all had very high sensitivity (> 96%) but also high FDR (28% – 55%); specificity, accuracy, and *F*_1_ measures were also somewhat suboptimal (Figure 3A). Among these six methods, LTR_FINDER demonstrated the best balance of performance across metrics followed by MGEScan3 (Figure 3A).

LTR_FINDER is one of the most popular LTR methods, however, it runs on a single CPU and is prohibitively slow for large genomes. We hypothesized that complete sequences of highly complex genomes may contain a large number of complicated nesting structures that cause the LTR_FINDER algorithm to iterate through loops. To break down these nesting structures, we developed a multithreading wrapper that cuts chromosome sequences into shorter segments and executes LTR_FINDER in parallel. Using this method, we see as much as 9,500X increase in speed for plant genomes varying from 120 Mb to 14.5 Gb (Table S3). For the 14.5 Gb bread wheat genome, the original LTR_FINDER took 10,169 hours, or 1.16 years, to complete, while the multithreading version completed in 72 minutes on a modern server with 36 CPUs, demonstrating an 8,500X increase in speed (Table S3). Among the genomes we tested, the parallel version of LTR_FINDER produced slightly different numbers of LTR candidates when compared to those generated using the original version (0% – 2.73%; Table S3), which is likely due to the use of the dynamic task control approach for processing heavily nested regions. Given the substantial speed improvement (Table S3) and high-quality results (Figure 3), we consider the parallel version to be a promising solution for large genomes. We used this parallel version of LTR_FINDER for all subsequent analyses in this study.

LTR_retriever does not have its own search engine; rather it was designed as a stringent filtering method for raw results of other LTR programs. LTR_retriever can process results of all six aforementioned LTR methods or any combination of them. We used LTR_retriever in conjunction with each of the six programs and with all six programs together to benchmark performance. Our results show that LTR_retriever has consistently high specificity (94.8% ± 3%), accuracy (92.2% ± 3%), precision (84.9% ± 7%), *F*_1_ measure (82.4% ± 10%) and low FDR (15.1% ± 7%) (Figure 3B; Table S1B). The sensitivity of LTR_retriever is also high (≥ 93%), except when used in combination with LTR_STRUC and LtrDetector (Figure 3B; Table S1B). This is due to the imprecisely defined sequence boundaries of LTR candidates of these two methods, preventing LTR_retriever from finding microstructures like TSD and terminal motifs [39], yielding a high false negative rate.

Overall, LTR_retriever presents the best compromise between sensitivity and specificity. LTR_retriever also generated the most compact LTR library in comparison to the other programs (Table S2), allowing efficient and precise whole-genome LTR annotations. It is not necessary to run all six structure-based programs along with LTR_retriever. Instead, the combination of LTR_FINDER and LTRharvest with LTR_retriever achieved the best performance and the shortest processing time as previously demonstrated (Table S2) [39].

### Comparison of non-LTR Annotators

Non-LTR retrotransposons include LINEs and SINEs that propagate via reverse transcription of RNA intermediates [16]. Identification of non-LTR retrotransposons is very challenging due to the lack of a long terminal repeat structure and because their sequences often degenerate relatively quickly [32]. In addition to the general repeat annotators described above, we also benchmarked a dedicated database for SINEs (SINEBase) and three structure-based methods.

SINEBase [32] is a species-agnostic database that performed similarly to the non-rice Repbase library (Figure 4B). Interestingly, the specialized structure-based annotation methods, including MGEScan3, SINE-Finder, and SINE_Scan also exhibited suboptimal sensitivity (< 60%) and very high FDRs (51% – 95%) (Figure 4; Table S1C). SINE_Scan is a successor of SINE-Finder, which aims to detect all known types of SINEs with higher accuracy [49]. Based on our results, SINE_Scan did have a much lower FDR compared to SINE-Finder; however, its sensitivity was also much lower (Figure 4B).

It is possible that SINEs are under annotated in the curated library, which may contribute to the high FDR values that were observed across programs. To test the validity of SINE candidates, we followed instructions in the SINE_Scan package and manually inspected head and tail alignments of all candidate SINE families (n = 35). Out of 35 candidate families, we found six that possess clear sequence boundaries with poly-A or poly-T tails and were longer than 99 bp. All of these six families were present in the curated library, indicating the high FDR is a product of false discovery rather than a limitation of the curated library being used to evaluate these programs.

In summary, we found general methods such as RepeatModeler, the non-rice Repbase, and SINEBase provided high-quality annotations for non-LTR retrotransposons, while structural based methods such as MGEScan3, SINE-Finder, and SINE_Scan have low sensitivity and high rates of false discovery. Therefore, researchers may want to use RepeatModeler for *de novo* annotation of non-LTR elements, and supplement these annotations with SINEBase or Repbase.

### Comparison of TIR Annotators

TIR transposons are a subclass of TEs that carry inverted repeats at both of their ends [16]. Miniature inverted transposable elements (MITEs) are a special kind of TIR transposon that lack any coding potential (non-autonomous) and are usually shorter than 600 bp [16]. These elements are highly abundant in eukaryotic genomes, and, as such, there are a large number of annotation programs designed to identify them. We tested P-MITE [31], a specialized database of curated plant MITEs, and IRF [50], TIR-Learner [17], and GRF (*grf-main -c 0*) (https://github.com/bioinfolabmu/GenericRepeatFinder), which structurally identify TIR elements, and finally MITE-Hunter [51], detectMITE [52], MUSTv2 [53], miteFinderII [54], MITE-Tracker [55], and GRF (*grf-mite*), which structurally identify MITEs specifically.

The P-MITE database performed similarly to what we observed for classifications from the general repeat annotators; the rice-specific database annotated TIR elements accurately and sensitively (P-MITE_rice), while the non-rice database (P-MITE_norice) had very low FDR and low sensitivity (Figure 5B), suggesting the necessity of using structure-based TIR and MITE detection methods for *de novo* annotation.

For TIR annotation using structure-based methods, IRF, GRF with educated parameters (GRF-TIR_edu), and TIR-Learner, all had high sensitivity (> 90%; Figure 5C; Table S1D), however, IRF and GRF-TIR_edu performed poorly for the remaining metrics (Figure 5C). The poor performance of these programs is due to the large number of candidates they identified, with 4.7 Gb and 630 Gb (13X – 1,684X the size of the 374 Mb rice genome) of raw TIR candidate sequences produced, respectively. The majority of raw candidate sequences were overlapping and nested within each other. The output of these programs was substantially filtered and condensed (Additional File 1; Table S2), and, despite this, still had poor performance based on our analysis metrics (Figure 5C). TIR-Learner demonstrates relatively high sensitivity, specificity, and accuracy (Figure 5C), which is promising for TIR annotation.

For structure-based MITE annotation, GRF with educated parameters (GRF-mite_edu) also produced large output files similar to IRF and GRF-TIR_edu. After filtering for false discovery and redundancy (Additional File 1), the candidate sequence file was reduced from 47 Gb (130X the size of the rice genome) to 10 Mb (Table S2). Still, given its inferior annotation performance relative to other MITE methods (Figure 5B), GRF-mite_edu is not ideal for *de novo* annotation. Interestingly, GRF with default parameters (GRF-mite_dft) had high performance, similar to MITE-Hunter and MITE-Tracker (Figure 5B). This is mostly due to changing the internal region length from default 780 bp to 10 kb (Additional File 1), which captured significantly more non-MITE sequences, suggesting the default parameters of GRF may have been optimized for MITE detection. These three MITE methods all had high specificity (≥ 95%) and accuracy (≥ 94%), reasonable sensitivity (79% – 84%), but somewhat lower precision (64% – 79%) (Figure 5B; Table S1D), suggesting high potentials for these programs. miteFinderII and detectMITE also had high performance but with comparatively lower sensitivity and lower specificity and accuracy, respectively (Figure 5B; Table S1D). MUSTv2 performed similar to GRF-mite_edu and worse than other MITE programs (Figure 5B).

We identified promising methods for TIR transposon and MITE annotation including TIR-Learner and MITE-Hunter, MITE-Tracker, and GRF-mite_dft, respectively. These methods all have high-performance metrics but somewhat high FDR (Figure 5), indicating each program generated annotations that covered our curated library as well as additional potential TEs. We are aware that our curated library is likely incomplete, and these new candidates could be real TIR elements or MITEs. We compared these new TE candidates with the curated library and to TIR element-specific conserved domains (Additional File 1). On an element basis, we found over half (4,104 out of 13,317 novel TIR elements and 11,885 out of 18,093 novel MITEs) of the elements shared similar TIR sequences with our curated library, but included variation in their internal sequences, with few elements showing potential to be autonomous (Figure 5D). Such variation is typical for nonautonomous TIR transposons, such as *Ds* elements [56]. For MITE candidates with novel TIRs, the majority had more than three copies in the rice genome (Figure 5D), suggesting these are likely novel TEs that were not included in the curated library. Out of the four MITE programs, MITE-Hunter identified sequences most similar to the curated library (Figure 5D).

TIR-Learner demonstrated great promise for structural annotation (Figure 5) and a large proportion of novel candidates were likely nonautonomous forms of known TIRs (Figure 5D). We found less than half of the novel TIR elements with novel TIRs had more than three copies in the rice genome (Figure 5D). This is because TIR candidates were not filtered based on copy number in TIR-Learner [17], given that some TEs may share similar TIRs but different internal regions (Figure 5D). Still, some of these could be contaminants such as LTR sequences. For example, we identified 6.38% of TIR-Learner candidates were actually LTR sequences using our curated library. After removal of these contaminants, the specificity and accuracy increased to 91.6% and 91.3%, respectively, while the sensitivity remained at ∼90%, and the FDR dropped from 57.3% to 30.8% (Figure 5C; Table S1D), suggesting that the high observed FDR was partially caused by misclassification of LTR sequences as TIR elements.

In summary, MITE-Hunter and TIR-Learner showed the highest performance for structural identification of MITEs and TIR elements (Figure 5BC), respectively, when TIR-Learner results were filtered to control false discovery (Figure 5C). RepeatModeler, Repbase, and P-MITE had high accuracy but low sensitivity (Figure 5AB), and could be used to supplement structural annotations of MITE and TIR elements.

### Comparison of *Helitron* Annotators

*Helitrons* are a subclass of DNA transposons that lack terminal repeats and do not generate target site duplications when transposed due to their rolling circle mechanism of transposition [57], making identification of these elements practically challenging. We found only one structure-based software, HelitronScanner [18], that is available, bug-free (no errors in our test), and produced *Helitron* predictions.

HelitronScanner produced 52 Mb of raw candidate sequences in rice (Table S2). Since *Helitrons* may capture DNA sequences when transposed, many non-*Helitron* TE sequences and even protein-coding sequences are present in the raw prediction. Nested insertions between different TE classes are also likely. Using the curated library, we annotated 1.8% of these *Helitron* candidates consisted of non-LTR sequences (LINEs and SINEs); 21% were LTR sequences and 11% were TIR sequences. With no filter applied, these *Helitron* candidates would include all classes of TEs, resulting in a high false discovery rate (93.7%; Table S1E) and low annotation performance (Figure 6A). To control for false discovery, we filtered captured sequences and candidates that lacked the *Helitron* signature 5’-TC…CTRR-3’ (R = G or A) terminal sequence structure, as well as those not inserted into AT or TT target sites (Additional File 1) [58]. We also removed non-*Helitron* TE sequences in these candidates using the curated library. After these filterings and cleaning, both the specificity and accuracy improved to 86%, while sensitivity was maintained at 95% (Figure 6A; Table S1E).

Similar to TIR-Learner for TIR element identification, HelitronScanner identified most of the curated *Helitrons* in the curated library, and also many additional elements not contained in the library (Figure 6A). We further filtered these candidates with the EDTA pipeline (Methods) and annotated the rice genome. Our filters yielded annotated sequences with similar mean and median length compared to the curated annotation, but this covered 13.9% of the rice genome compared to 3.6% from annotation using the curated library (Figure 6B). Evaluation of the 30 bp sequences of both terminals with 10 bp flanking sequences as sequence logos showed the AT or TT target sites we required in our filtering, and also that these candidates clearly have the canonical terminal structure 5’-TC…CTRR-3’ (with 5’-TC…CTAG-3’ dominating) which is required by HelitronScanner (Figure 6C). The candidates were also located in relatively AT-rich regions with significantly higher AT content in the 5’ terminal, consistent with previous observations by Yang and Bennetzen (2009) [59]. We found enriched CG content at the 3’ terminals especially at the −13 and −14 positions, which could produce a hairpin loop, a canonical *Helitron* feature [18]. While these elements contain the terminal features of a *Helitron*, this does not confirm the validity of these elements.

Overall, the filtered HelitronScanner candidates covered the majority of *Helitrons* in the curated library and identified a large number of additional elements that displayed a number of canonical *Helitron* features. Based on the EDTA filtered test library, about 14% of the genomic sequence of rice was annotated as *Helitron*-like (Figure 6B), which could be due to other DNA sequences being captured in *Helitrons*. To further confirm this result, meticulous curation and intra-specific comparisons are needed [18, 58].

### Comparison of Resource Consumption and Usage

In this study, we benchmarked 25 TE annotation programs and three databases, while nine others were attempted with failure due to a variety of reasons: 1. Lack of maintenance with unresolved bugs; 2. Outdated programs required by the software and a lack of alternatives; 3. Required programs or databases are not open-source; and 4. Programs take too long to run. For programs that were run successfully, some were more challenging than others. One of the main obstacles was installation. We found compile-free and precompiled programs were the easiest to use, followed by those available via conda and bioconda [60].

In addition to benchmarking the quality of the output of each program, we also benchmarked the algorithmic efficiency of these TE annotation programs. Since these programs were executed in different high-performance computational platforms (Table S2), algorithmic performance could be slightly variable. Overall, most programs completed within 24 hours with an average of 5.5 hours (Table S2). Longer run time was not associated with higher performance in terms of the six analysis metrics, and could become a barrier for annotation of large genomes. Most programs were not memory intensive, with a minimum of 7.2 Mbyte (SINE-Finder), an average of 8.7 Gbyte, and a maximum of 76 Gbyte (the GRF-LTR_FINDER method) (Table S2). Approximately two-thirds of the programs can be multi-threaded. However, the average CPU usage of programs was not significantly correlated with run time (*r* = −0.19, *p* = 0.26, *F*-test), indicating run time is primarily determined by algorithmic efficiency.

### Construction and Benchmarking of the EDTA pipeline

From the benchmarking results, we identified a set of programs that presented high sensitivity, specificity, and accuracy, but, in some instances, high FDR. Using these programs, we have developed a pipeline called <u>E</u>xtensive *<u>d</u>e-novo* <u>T</u>E <u>A</u>nnotator (EDTA), which combines the best-performing programs and subsequent filtering methods for *de novo* identification of each TE class and compiles the results into a comprehensive TE library. The EDTA pipeline incorporates LTRharvest, the parallel version of LTR_FINDER, LTR_retriever, TIR-Learner, MITE-Hunter, HelitronScanner, and RepeatModeler as well as customized filtering scripts (Figure 7A). Basic filters for LTR candidates, TIR candidates, MITE candidates, and *Helitron* candidates were the same as those applied for benchmarking of programs in previous sections, and control minimum sequence length and remove tandem repeats (Stage 0; Methods). Advanced filters use high-quality TEs identified in stage 0 candidates to remove misclassified sequences (Stage 1; Methods).

**Figure 7.**
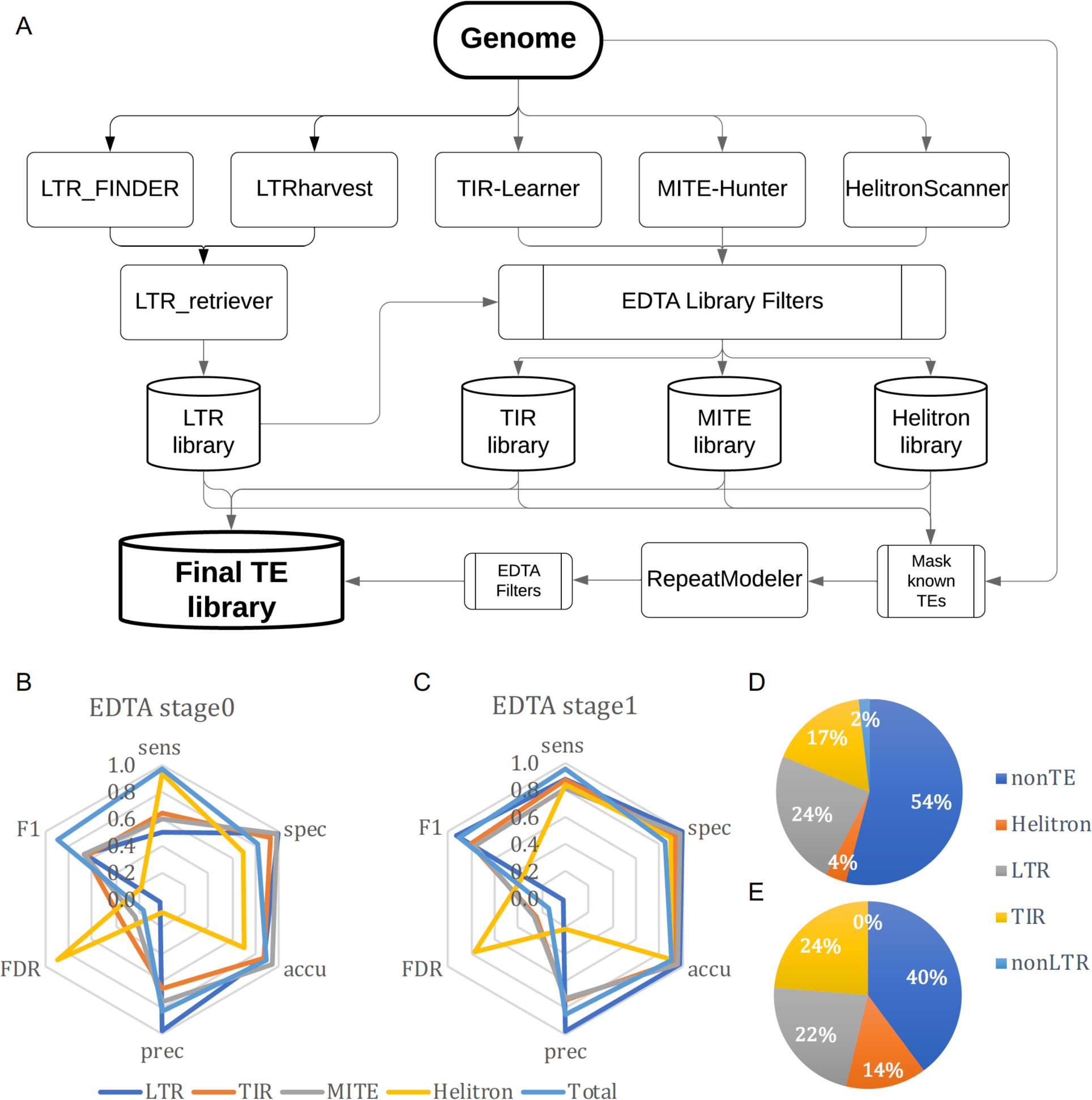
The Extensive *de-novo* TE Annotator (EDTA) pipeline. (A) The EDTA workflow. LTR retrotransposons, TIR elements, MITEs, and *Helitron* candidates are identified from the genome sequence. Sub-libraries (such as LTR library, TIR library, etc) are filtered using EDTA library filtering scripts (including both basic filters and advanced filters, see Methods for details) for removal of misclassified TEs, and are then used to mask TEs in the genome. The unmasked part of the genome is processed by RepeatModeler to identify non-LTR retrotransposons and any unclassified TEs that are missed by the structure-based library. Nested insertions and protein coding sequences are removed in the final step to generate the final TE library. Performance of (B) EDTA stage 0 sublibraries and (C) EDTA stage 1 sublibraries after basic filterings and advanced filterings, respectively. Annotation of the rice genome using (D) the curated library and (E) the final EDTA generated library.

To test the performance of the EDTA pipeline, we annotated the rice genome using the curated TE library and the test library generated from the EDTA pipeline. Performance metrics of stage 0 results showed very low sensitivity (< 65%) for annotation of LTR elements, TIR elements, and MITEs, and also suboptimal specificity (∼70%) and accuracy (∼70%) for *Helitron* annotations (Figure 7B; Table S1F). This is due to the nested TEs, captured TEs, or false discovery in *Helitron* candidates that impair the annotation performance in the combined stage 0 library. After reciprocal removal of misclassified TEs in each category (Stage 1; Figure 7A; Methods), the performance metrics were high for the final EDTA pipeline annotation (Figure 7C). For all four TE subclasses and the overall repetitive sequences, the annotation sensitivity ranged from 81% – 96%, specificity ranged from 85% – 99%, and accuracy ranged from 89% – 97% (Table S1F). FDRs of these categories ranged from 2% – 27%, with the exception of *Helitrons* that had 77% of the annotations not identified by the curated library (Table S1F). Our results from the benchmarking and the final EDTA pipeline indicate the potential need for more thorough *Helitron* annotation using our pipeline (Figure 6). However, careful verification and curation are still necessary to further confirm this result.

Using the EDTA pipeline, we could not identify non-LTR retrotransposons using the RepeatModeler module (Figure 7E). This is likely due to the low level of non-LTR elements in the rice genome (Table 1; Figure 7D) that could have been misclassified as other TE subclasses. Further annotation of non-LTR retrotransposons is necessary to exhaustively annotate TEs in the genome. As new programs become available for non-LTR elements, they will be benchmarked and potentially added to the EDTA pipeline based on performance metrics.

## Discussion

Recent innovations in third-generation (*i.e.*, long read) sequencing have enabled rapid and high-quality assembly of the repetitive fraction of genomes, creating an opportunity and need for high-throughput annotation of transposable elements. Annotation of TEs presents a substantial algorithmic and computational challenge. Different classes of TEs have distinct sequence characteristics, which has led to the development of software programs for each type. While anecdotally researchers have known the strengths and weaknesses of each of these methods, no comprehensive benchmarking study has quantified their relative annotation (*i.e.*, sensitivity, specificity, etc.) and computational (*i.e.*, run time, memory requirements, etc.) metrics. We have exhaustively tested these programs against a high-quality, manually curated rice TE library and have compiled the best-performing software as part of a comprehensive TE annotation pipeline known as EDTA.

All TEs are or were capable of transposition in the genome. However, the ability to amplify varies dramatically among different TE families. In fact, only a few TE families can amplify to high copy number. For example, in maize, the top 20 families of LTR retrotransposons comprise ∼70% of the genome, whereas the remainder (380 or more) comprise only ∼5% [61]. From this perspective, if a TE identification program captures elements with high copy number, the majority of the TE body in the genome will be characterized. Consistent with this notion, we observed that all general repeat identification programs, which depend on repetitive target sequences, performed well (high sensitivity and specificity, good precision and accuracy; Figure 2A). Most importantly, the results from these programs are associated with very low FDR, suggesting when a sequence is repetitive to a certain degree, it is very likely to be a TE. However, most repeats from general programs are not classified and their sequence boundaries are often approximate. Not all tasks require this information. For example, repetitive sequences are usually masked prior to gene annotation to minimize interference. For such purposes, general repeat identification programs and subsequent filtering for duplicated genes would suffice.

In contrast to the general repeat annotators, structure-based programs can identify low- or even single-copy elements and are therefore complementary. Moreover, these programs provide the exact coordinates of elements and are ideal for targeted study of TEs and their interactions with other components in the genome. However, based on our results, the majority of structure-based programs are associated with high FDR (up to 95%), and such error could be propagated in subsequent analyses. One factor contributing to this high error rate is misidentification due to nested insertion of TEs from different classes. We have developed an approach to minimize this issue by cross-checking sequences derived from programs for different classes of TEs. Another potential strategy to reduce FDR is to incorporate copy number control but this would actually compromise the most important advantage of structure-based programs. Thus, this is an unsolvable problem without improvement to structure-based programs; particularly those for non-LTR retrotransposons and *Helitrons*. While more specific search engines or efficient filters may reduce the FDR, some level of manual curation may still be necessary for the generation of high-quality libraries.

Few species beyond rice have TE libraries of sufficient quality and genomes that are tractable enough to be used for benchmarking purposes. Furthermore, TEs comprise a relatively high proportion of the rice genome (∼47%), and our extensive manual curation efforts make it one of the only species in which a benchmarking study can reliably calculate true positive, false positive, true negative, and false negative rates across annotation programs. However, relative performance of TE annotation programs should be similar across systems. Programs have primarily been developed to detect specific types of TEs and are largely agnostic to species. This is possible because classes of TEs generally have similar structures across species [14,16,18]. Throughout this benchmarking exercise, we have based our tuning of programs (*i.e.*, our educated parameters) on current knowledge of the structure of each target TE class [14,16,18], which, again, is not specialized to a particular system. As an example of the broad utility of these methods, the LTR_retriever program [39] has been tested for annotation of Arabidopsis, rice, Maize, and sacred lotus (*Nelumbo nucifera*) [62], and demonstrated similar performance across systems.

We do anticipate some limits to the broad applicability of the EDTA pipeline across systems. For instance, TIR-Learner uses machine learning to train TIR classifiers, and we have used the curated rice TIR element-trained classifier in this study. The method could be made more general by using a broader set of TIR transposons to train the classifier, such as curated elements in the P-MITE database [31]. Furthermore, based on our metrics, the performance of methods for detecting the non-LTR elements (*i.e.*, SINEs and LINEs) was generally suboptimal and better algorithms are needed. Particularly, there is no structure-based program available for the identification of LINEs. The EDTA package may therefore miss a number of elements in, for instance, vertebrate genomes that contain many SINEs and LINEs [63]. Finally, our knowledge of TE structure is rapidly expanding, and parameterization and tuning of methods will therefore need to be continually updated. For example, variation in terminal motifs and target site duplication in LTR elements was previously poorly characterized. In the development of LTR_retriever, it was found that the terminal motif 5’-TG..CA-3’ occurs 99% of the time and that the vast majority of LTR TSDs are 5bp [39]. While some programs set very flexible parameters for these features (*e.g.*, LTRharvest), in our implementation of LTR_retriever we applied our new knowledge during curation and observed a substantial improvement in performance.

Moving forward, we see opportunities for improved annotation of highly variable TE classes including MITE/TIR elements and SINE/LINE, where, upon insertion, mutations and indels can be created. In these situations, construction of a consensus sequence is necessary for more precise TE annotation. Many programs do not currently have this feature. The GRF program for detection of interspersed repeats (*grf-intersperse*) has a consensus function, but the program does not allow indels, resulting in the lowest sensitivity but also the lowest FDR. For SINE/LINE detection, we found very low sensitivity and very high FDR, which is likely due to variation in these TEs (*e.g.*, most LINEs are truncated upon insertion) and the lack of terminal repeats, making detection very challenging. Further development of consensus-based methods will be important. As new methods are generated and existing methods are improved, they will be benchmarked relative to our rice library and included in the EDTA pipeline when they result in a marked increase in annotation performance.

## Conclusions

Advances in sequencing technology are facilitating assembly of the repetitive portion of the genome, which necessitates the annotation of these features. Using a highly curated library of rice TEs, we have created a benchmarking platform to test TE annotation software. We used this platform to exhaustively test currently available software based on output (*i.e.*, sensitivity, specificity, etc.) as well as the performance of the software (*i.e.*, run time, memory usage, etc.). From this benchmarking exercise, the EDTA pipeline was developed that combines the highest performing software with necessary filtering and processing scripts such that the pipeline can be applied to any new genome assembly.

## Methods

### Manual curation of transposable elements in rice

Manual curation of TEs in rice was started after the release of the map-based rice genome in 2005 [22]. Repetitive sequences in the rice genome were compiled by RECON [43] with a copy number cutoff of 10. Details for manual curation of LTR sequences were previously described in the LTR_retriever paper [39]. In brief, for the curation of LTR retrotransposons, we first collected known LTR elements and used them to mask LTR candidates. Unmasked candidates were manually checked for terminal motifs, TSD sequences, and conserved coding sequences. Terminal repeats were aligned with extended sequences, from which candidates were discarded if alignments extended beyond their boundaries. For the curation of non-LTR retrotransposons, new candidates were required to have a poly-A tail and TSD. We also collected 13 curated SINE elements from [49] to complement our library.

For curation of DNA TEs with TIRs, flanking sequences (100 bp or longer, if necessary) were extracted and aligned using DIALIGN2 [64] to determine element boundaries. A boundary was defined as the position to which sequence homology is conserved over more than half of the aligned sequences. Then, sequences with defined boundaries were manually examined for the presence of TSD. To classify the TEs into families, features in the terminal and TSD sequences were used. Each transposon family is associated with distinct features in their terminal sequences and TSDs, which can be used to identify and classify elements into their respective families [14]. For *Helitron*s, each representative sequence requires at least two copies with intact terminal sequences, distinct flanking sequences, and inserts into “AT” target sites.

To make our non-redundant curated library, each new TE candidate was first masked by the current library. The unmasked candidates were further checked for structural integrity and conserved domains. For candidates that were partially masked and presented as true elements, the “80-80-80” rule (≥ 80% of the query aligned with ≥ 80% of identity and the alignment is ≥ 80 bp long) was applied to determine whether this element would be retained. For elements containing detectable known nested insertions, the nested portions were removed and the remaining regions were joined as a sequence. Finally, protein-coding sequences were removed using the ProtExcluder package [65]. The curated library version 6.9.5 was used in this study and is available as part of the EDTA toolkit.

### Calculation of benchmarking metrics

The curated TE annotation of the rice genome (*Oryza sativa* L. ssp. *japonica* cv. ’Nipponbare’ v. MSU7) was created using the standard library (v6.9.5) and RepeatMasker v4.0.8 with parameters “-pa 36 -q -no_is -norna -nolow -div 40 -cutoff 225”. These parameters identified homologous sequences with up to 40% divergence without detecting bacterial insertion elements, small RNA (pseudo) genes, and low complexity DNA. This annotation was used as the curated annotation for the calculation of benchmarking metrics. For genomic regions that cover more than 80% of a TE sequence in the curated library, the region was counted as a complete copy, and those that covered less than 80% were counted as a fragmented copy.

When we obtained a non-redundant test library from a target program (details in the next section), the test library was used to annotate the rice genome with the same RepeatMasker parameters, except that the test library was provided as a custom library. Then, the testing annotation was compared to the curated annotation for calculations of sensitivity, specificity, accuracy, precision, FDR, and *F*_1_ measures (Figure 1). These six metrics were calculated using the script “lib-test.pl” in our EDTA toolkit.

### Execution of TE programs

We exhaustively searched the literature for open-source programs and databases that have been developed for both general repeat annotation and structural annotation. We executed each of these programs to obtain candidate sequences or downloaded sequences from specialized databases. All programs were executed using parameters consistent with current knowledge of TE structure (educated parameters). A description of each of these programs, observations we made about accessibility/ease of use of these programs, and the specific parameter options that were used are provided in Additional File 1. To benchmark the algorithmic efficiency, these programs were executed in multiple high-performance computing platforms (Table S2). Run time (wall clock), average CPU usage, and maximum memory consumption were recorded using “/usr/bin/time -v”.

After we obtained raw sequences from programs, we went through three steps to construct non-redundant test libraries. The first step was to remove short tandem repeat contamination sequences that were present in the raw candidates. Identification of tandem sequences was achieved by Tandem Repeats Finder [66] with parameters “2 7 7 80 10 3000 2000 -ngs -h -l 6”. The second step was to remove missing characters (Ns) in candidates as well as short sequences. The minimum sequence length was set to 80 bp for TIR candidates and 100 bp for other types of TE candidates. We used the script “cleanup_tandem.pl” in the LTR_retriever package [39] for the first two steps with parameters “-misschar N -nc 50000 –nr 0.9 -minlen 100 (or 80) -minscore 3000 -trf 1 -cleanN 1”. The third step was to remove redundant sequences and nested insertions, which was achieved using the script “cleanup_nested.pl” in the LTR_retriever package [39] with default parameters. The third step was iterated five times to resolve heavily nested TEs for a thorough reduction of sequence redundancy. The resulting sequences were used as the non-redundant test library for the focal programs. Databases were used directly as test libraries without any filterings or manipulations.

### Construction of the Extensive *de novo* TE Annotator pipeline

Extensive *de-novo* TE Annotator (EDTA) is a pipeline for comprehensive and high-quality TE annotation for newly sequenced eukaryotic genomes. We combined open-source programs that are either specialized for a particular subclass of TEs or general for all repetitive sequences. The programs we selected had the highest performance from our benchmarking and together deliver the best TE annotation for a new genome that is possible given current program performance. Still, based on our benchmarking results, substantial contamination will exist due to misclassification of elements, nested insertions, and sequences captured by TEs.

The EDTA pipeline contains a set of scripts for filtering the output of each program to reduce the overall false discovery rate. The first set of scripts included in EDTA apply a basic filter for each of the initial predictions to remove tandem repeats and short sequences (< 80 bp for TIR elements and < 100 bp for LTR elements and *Helitrons*). For LTR candidates identified by LTRharvest and LTR_FINDER, false discoveries are filtered by LTR_retriever. For TIR candidates identified by TIR-Learner, sequences are reclassified as MITEs if their length is ≤ 600 bp. MITE-Hunter results are masked using TIR-Learner results to remove duplicated candidates. Unmasked MITE candidates are added to the TIR-Learner results to supplement MITE discovery. For *Helitron* candidates reported by HelitronScanner, filters based on target site (AT or TT) and prediction scores (≥ 12) are performed (Additional File 1). After these basic filtering steps, TE candidates are named stage 0.

Advanced filters are necessary to generate a comprehensive and high-quality TE library. First, we identified contaminant-like sequences in stage 0 LTR and TIR sublibraries using LTRharvest, MITE-Hunter, and HelitronScanner with default parameters. In particular, LTRharvest is used to identify LTR-like sequences in the TIR sublibrary as suggested by Figure 5C; MITE-Hunter is used to identify MITE-like sequences in the LTR sublibrary as suggested in [39]; HelitronScanner is used to identify *Helitron*-like sequences in LTR and TIR sublibraries. Then stage 0 sublibraries are masked and cleaned using these contaminant-like sequences to generate high-quality TE sequences (stg0.HQ). As solo LTRs are the most prevalent form of LTR contamination, only LTR regions from the stg0.HQ LTR elements are used to clean up LTR contamination in TIR and *Helitron* candidates. In particular, stg0.HQ.LTR sequences are used to mask and clean the stage 0 TIR sublibrary; and stg0.HQ.LTR and stg0.HQ TIR sequences are used to mask and clean the stage 0 *Helitron* sublibrary.

After these reciprocal filtering steps (Figure 7), updated sublibraries are aggregated and subjected to three rounds of nested insertion removal and clustering, which generates the non-redundant stage 1 library. Because LTR_retriever serves as a strong filter of results from LTRharvest and LTR_FINDER, no further filtering was necessary (LTR.stage0 = LTR.stage1). Non-redundant stage 1 TEs are then used to mask the genome. The remaining unmasked portion of the genome is scanned by RepeatModeler with default parameters to identify non-LTR retrotransposons and any unclassified TEs that are missed by structure-based TE identification. Finally, all remaining TEs are aggregated, protein-coding sequences are filtered, and two more rounds of redundancy reduction are completed in order to produce the final TE library. This library has RepeatMasker-readable sequence names and can be used to annotate whole-genome TE sequences.

## List of abbreviations

TE: Transposable Elements
LTR: Long Terminal Repeat
LINE: Long Interspersed Nuclear Element
SINE: Short Interspersed Nuclear Element
MITE: Miniature Inverted Transposable Element
TIR: Terminal Inverted Repeat
TSD: Target Site Duplication
TP: True Positives
FP: False Positives
TN: True Negative
FN: False Negatives
GRF: Generic Repeat Finder
EDTA: Extensive *de-novo* TE Annotator

## Declarations

### Ethics approval and consent to participate

Not Applicable

### Consent for publication

Not Applicable

### Availability of data and materials

The curated rice library and all scripts are freely available at https://github.com/oushujun/EDTA.

## Author’s contributions

SO, MBH, and CNH conceived the study. SO, WS, YL, KC, NJ conducted the analyses. SO developed the EDTA package. SO, MBH, CNH, and NJ wrote the manuscript. All authors read and approved the final manuscript.

## Acknowledgements

We wish to acknowledge Margaret Woodhouse and Jeffrey Ross-Ibarra for helpful feedback on a previous version of this manuscript.

## Additional Files

**Additional File 1.** Supplementary Methods.

**Additional File 2.** Supplementary Tables 1-3. Supplemental Table 1 contains a summary of benchmarking results. Supplemental Table 2 contains a summary of time and resource consumption for each program. Supplemental Table 3 contains benchmarking information for the parallel wrapper of LTR_FINDER.

