## Additional File 1 for "Benchmarking Transposable Element Annotation Methods for Creation of a Streamlined, Comprehensive Pipeline"

### Additional File 1. Supplementary Methods

#### Parameter settings of TE programs

Details regarding program parameters and user experience are described below. For processing of results, please refer to the Methods section of this study.

*General repeat annotators*

**RECON** [[1]](https://paperpile.com/c/Ovk0Hl/xhYU) was developed for *de novo* identification and classification of repeat sequences based on multiple sequence alignment information. We used the RepeatModeler patched version (v1.08) in our test. The initial all-vs-all pairwise comparison of the rice genome sequence was performed with BLAST (v2.2.31). Removal of self-alignments and formatting for RECON required using the script “Filter_and_format_blast_for_RECON.pl” in our Extensive *de-novo* TE Annotator (EDTA) toolkit. Because RECON is a single-threaded program, we ran RECON separately for each chromosome with default parameters, and combined the results afterwards.

**RepeatScout** [[2]](https://paperpile.com/c/Ovk0Hl/PObp) is a *k*-mer based program to identify repetitive sequences. The version 1.0.5 was used in this study, which is single-threaded. We used 16 bp as the starting seed length (-l 16) and minimum copy number of 10 (--thresh=10) for *de novo* repeat identification.

**RepeatModeler** [[3]](https://paperpile.com/c/Ovk0Hl/8ZJO) is a *de novo* repeat identification and classification program which was built based on RepeatScout [[2]](https://paperpile.com/c/Ovk0Hl/PObp) and RECON [[1]](https://paperpile.com/c/Ovk0Hl/xhYU). RepeatModeler helps to refine TE boundaries, classify TEs based on a genomic database, and construct non-redundant TE libraries for whole-genome annotation. We used the v1.0.11 version of RepeatModeler that was multithreading-enabled with the NCBI BLAST engine (-engine ncbi). We ran RepeatModeler with default parameters.

**Red** [[4]](https://paperpile.com/c/Ovk0Hl/1SNe) is a C++ program which enables rapid detection of repetitive sequences. The software is not dependent on other programs and is available in bioconda. Parameters for running Red are automatically defined based on the input genome sequence. Repeat candidates detected by Red are formatted as a list of genome coordinates; sequences were extracted using the “call_seq_by_list.pl” script in our EDTA toolkit.

**Generic Repeat Finder (GRF)** (<https://github.com/bioinfolabmu/GenericRepeatFinder>) is a C++ program that can detect multiple types of repeats including interspersed repeats, terminal inverted repeats (TIR, including MITEs), and terminal direct repeats (TDR, including LTR retrotransposons). The grf-intersperse module of GRF was used to detect interspersed repeats in groups (-f 0 -c 5), and the grf-alignment2 module was further used to generate consensus sequences. The “get_interspersed_consensus.pl” script in our EDTA toolkit was developed to extract consensus sequences with filtering of the minimum sequence length of 80 bp. The CD-HIT [[5]](https://paperpile.com/c/Ovk0Hl/0a2b) program was used to further cluster consensus sequences with a minimum of 99% coverage and 80% identity (-c 0.80 -n 5 -d 0 -aL 0.99 -s 0.8 -M 0).

**RepBase** [[6]](https://paperpile.com/c/Ovk0Hl/w23c) is a database of consensus TE sequences derived from known eukaryotic genomes, which has been incorporated in RepeatMasker [[3]](https://paperpile.com/c/Ovk0Hl/8ZJO) for whole-genome TE annotations. We used the rice database in RepBase to benchmark its annotation performance by comparing to our curated database. The RepeatMasker version v4.0.8 and RepBase version 20170127 was used (-species rice -e Crossmatch). We also tested the non-rice database in RepBase for *de novo* TE annotation of rice. All core and comprehensive sequences that are not derived from rice were used in the test.

Other methods that were tried included REPET [[7]](https://paperpile.com/c/Ovk0Hl/vROr), CENSOR [[8]](https://paperpile.com/c/Ovk0Hl/PJZJ), and PILER [[9]](https://paperpile.com/c/Ovk0Hl/uMnX). CENSOR requires a .map file that was not described in the manual and on the website, thus we could not decipher its usage. REPET and CENSOR rely on the RepBase library, which has been commercialized recently and requires membership for any use and is not compatible with the open-source toolkit we developed. PILER relies on whole-genome self-alignment for detection of TEs. The step uses PALS [[9]](https://paperpile.com/c/Ovk0Hl/uMnX), which has been deprecated.

*LTR retrotransposons*

**LTR_STRUC** [[10]](https://paperpile.com/c/Ovk0Hl/SxBQ) is one of the earliest programs developed for detection of long terminal repeat (LTR) retrotransposons. It is Windows-based and requires a non-commercial license. LTR_STRUC requires no parameter settings and generates four files for each candidate. The script “convert_ltr_struc.pl” in the LTR_retriever program [[11]](https://paperpile.com/c/Ovk0Hl/CT05) was used to extract coordinate information from LTR_STRUC outputs.

**LTR_FINDER** [[12]](https://paperpile.com/c/Ovk0Hl/nwsc) is a commonly used program for *de novo* identification of LTR retrotransposons. It allows many parameters to be set. However, LTR_FINDER only allows one-CPU for all jobs, limiting the scalability of this program. To work around this limitation, we developed a multithreading wrapper called “LTR_FINDER_parallel.pl” in our EDTA toolkit, which will chop the genome into small pieces to run LTR_FINDER in parallel. We used 1Mb chunks and 300 timeout seconds for each chunk (-harvest_out -size 1000000 -time 300). LTR_FINDER version 1.0.6 was used with the minimum length of LTR regions, the maximum length of LTR regions, the maximum length of the whole candidate, and the maximum divergence between terminal repeats set to 100 bp, 7000 bp, 15,000 bp, and 85%, respectively (-w 2 -C -D 15000 -d 1000 -L 7000 -l 100 -p 20 -M 0.85). The script “convert_ltr_finder.pl” in the LTR_retriever program [[11]](https://paperpile.com/c/Ovk0Hl/CT05) was used to extract coordinate information from LTR_FINDER outputs.

We used four genomes to benchmark the performance of the original and the parallel version of LTR_FINDER. These four genomes are *Arabidopsis thaliana* (v. TAIR10), rice (*Oryza sativa* v. MSU7), maize (*Zea mays* v. AGPv4), and bread wheat (*Tricitum* *aestivum* v. CS1.0). All these tests were run with the entire genome with parameters described above except for the wheat genome where the original LTR_FINDER was run on each chromosome and combined after completion.

**LTRharvest** [[13]](https://paperpile.com/c/Ovk0Hl/Ch9e) is another popular program for *de novo* LTR detection. The availability of precompiled binary code makes it user friendly. While LTRharvest runs on only 1 CPU, it has a relatively short run time. We used version 1.5.10 for our analyses, with parameters similar to what we applied in LTR_FINDER (-minlenltr 100 -maxlenltr 7000 -mintsd 4 -maxtsd 6 -similar 85 -vic 10 -seed 20 -seqids yes). We also specified the canonical terminal motif 5'-TG...CA-3' with a maximum of 1 bp mismatch for the detection of LTR candidates (-motif TGCA -motifmis 1).

**MGEScan3** [[14]](https://paperpile.com/c/Ovk0Hl/sED5) is a Galaxy-based program that combines MGEScan_LTR [[15]](https://paperpile.com/c/Ovk0Hl/I4QL) and MGEScan-non-LTR [[16]](https://paperpile.com/c/Ovk0Hl/ZPaW). Thus, it detects both LTR and non-LTR retrotransposons. The command line tool is easy to use, but does not allow specific parameters to be set for either algorithm. The program is multithreading-enabled and is relatively quick. The script “convert_MGEScan3.0.pl” in the LTR_retriever program [[11]](https://paperpile.com/c/Ovk0Hl/CT05) was used to extract coordinate information from MGEScan3 outputs.

**LtrDetector** (<https://github.com/TulsaBioinformaticsToolsmith/LtrDetector>) is a recently developed tool for *de novo* LTR annotation. Installation of this C++ program was straight-forward, though it requires a very new compiler (gcc/8.2.0). The default options for the LtrDetector parameters were used. The default bed format output of LtrDetector includes target site duplication (TSD) as part of the candidate. We used the script “convert_ltrdetector.pl” in the LTR_retriever program [[11]](https://paperpile.com/c/Ovk0Hl/CT05) to remove TSD coordinates and extract LTR information for further analyses.

**GRF** (<https://github.com/bioinfolabmu/GenericRepeatFinder>) can detect terminal direct repeats (TDR), which could be provided to a modified version of LTR_FINDER for detection of LTR retrotransposons. We first used our best practice parameters to obtain raw TDRs; however, the modified LTR_FINDER could not detect any LTR candidates. We then used the parameters provided in the manual (-c 2 --min_tr 100 --min_space 1090 --max_space 23490 --match 2 --mismatch 2 --indel 3 -f 1 -p 30) to run *grf-main*, and used *grf-filter* to filter out extreme TDRs (*grf-filter* 100 3500 1000 20000). Finally, the modified LTR_FINDER provided in the GRF package was executed following the options suggested in the program manual. Using these options, a large number of candidates were obtained, and coordinate information was extracted using the script “convert_ltr_finder.pl” in the LTR_retriever program [[11]](https://paperpile.com/c/Ovk0Hl/CT05) for further analysis.

**LTR_retriever** [[11]](https://paperpile.com/c/Ovk0Hl/CT05) is a Perl program that was designed to filter excessive false positives in *de novo* LTR detection and construct a high-quality non-redundant library for whole-genome annotation. Input files are very flexible for LTR_retriever; for example, it can utilize the output of LTR-related programs and produce high-quality LTR candidates. We ran LTR_retriever on outputs of LTR_STRUC, MGEScan3, LTR_FINDER, LTRharvest, LtrDetector, and GRF-LTR_FINDER, respectively, and also on all six program outputs combined. Since LTR_retriever has been optimized for the best balance between sensitivity and specificity, we used default parameters for all LTR_retriever analyses.

We also tried LTR_MINER [[17]](https://paperpile.com/c/Ovk0Hl/cN8z) and LTR Annotator [[18]](https://paperpile.com/c/Ovk0Hl/ECgZ). While LTR_MINER is a Perl program designed to annotate intact and solo LTR elements from RepeatMasker output files, we could not get it to work due to changes in the RepeatMasker output format in newer versions of the program. For LTR Annotator, we could not find any installation package associated with this name from both their paper and a Google search.

*nonLTR (SINE and LINE)*

**SINEBase** [[19]](https://paperpile.com/c/Ovk0Hl/tiOB) is a database of short interspersed nuclear elements (SINEs). We downloaded consensus sequences from <http://sines.eimb.ru/> in both SINEBank and LINEBank for a collection of SINE families and partner LINEs, respectively. A total of 228 elements were downloaded.

**MGEScan3** [[14]](https://paperpile.com/c/Ovk0Hl/sED5) was used to *de novo* detect non-LTR retrotransposons. There is no parameter settings for this program other than being multi-threadible. We found only a small number of candidates (20) using MGEScan3 in the rice genome.

**SINE-Finder** [[20]](https://paperpile.com/c/Ovk0Hl/t2OV) is a Python program designed to identify tRNA-derived SINEs. There is no installation required for this program and parameter settings are straightforward. However, only the forward strand of the input sequences could be searched without errors, which requires manually generating and providing the reverse complement strand for the complete search. A total of 431 raw candidates were identified.

**SINE_Scan** [[21]](https://paperpile.com/c/Ovk0Hl/qQFq) is a Perl program representing the latest development of *de novo* SINE identification methods which is based on SINE-Finder. SINE_Scan can identify all three known types of SINEs, which are tRNA, 7SLRNA, and 5SRNA. SINE_Scan allows users to start their analyses from different steps and generates multi-sequence alignment (MSA) files for each candidate for manual curation. We used default parameters to run SINE_Scan and identified a total of 35 SINE family candidates in rice.

*TIR*

**P-MITE** [[22]](https://paperpile.com/c/Ovk0Hl/dsR1) is a database of plant Miniature Inverted Transposable Elements (MITEs), which contains 3,527 MITE families obtained from 41 plant species. Download links for the P-MITE database were broken (<http://pmite.hzau.edu.cn/download/>). We obtained non-redundant and TSD-removed MITE sequences from Dr. Jiongjiong Chen. We used rice and non-rice datasets to test the annotation performance of the P-MITE database.

**IRF** [[23]](https://paperpile.com/c/Ovk0Hl/5STs) is a multi-platform software designed to identify inverted repeat elements. It has five required parameters and 19 optional parameters. Some of the IRF parameters are quite technical for inexperienced users, such as matching/mismatching scores, indel penalty, and match/indel probabilities. We used parameters of matching weight +2, mismatch penalty -3, indel penalty -5, default match and indel probabilities 80 and 10, respectively, minimum score of 20, maximum stem length of 10000, maximum loop of 10000, and identity value of 80 ( “2 3 5 80 10 20 10000 10000 -h -r 80 -t5 10000”). The rice genome was split into 12 chromosomes, and each chromosome was processed separately. The output of IRF software is a .dat file, with coordinates and other information, therefore, users need to take one more step to obtain the FASTA file from the IRF output.

**MITE-Hunter** [[24]](https://paperpile.com/c/Ovk0Hl/Aa1t) is a Perl program and installation requires formatdb, blastall, mdust, and muscle as prerequisites. The software blastall and formatdb are outdated, and have been replaced by the software package BLAST+. MITE-Hunter has 17 parameter options and some are redundant. We used parameters of maximum unmatched 2 bp in TIR region, the maximum length of 1000 bp, a minimum shared length of 80 bp between TEs that will be grouped together, and minimum copy number of 1 ( “-l 2 -w 1000 -L 80 -m 1 -S 12345678 -c 16”). One of the best features for MITE-Hunter is that this software can take checkpoints. The -S parameter allows the user to start the program from a specific step. Therefore, if there is anything wrong in the running process, users do not have to restart from the beginning. MITE-Hunter generates hundreds of intermediate files in the same folder, some of which are very large. This could cause difficulties for users to obtain the final output and organize files in the target folder.

**detectMITE** [[25]](https://paperpile.com/c/Ovk0Hl/UxsV) is an open-source MATLAB program for *de novo* detection of MITEs. Its installation requires a third-party software CD-HIT [[5]](https://paperpile.com/c/Ovk0Hl/0a2b). The output of detectMITE is a FASTA file that includes coordinates, TIR length, and TSD length. We identified MITEs of maximum length 1000 bp (matlab -nodisplay -nosplash -r tic; do_MITE_detection('Rice.fa', '-mite_maximum_length',1000, '-genome','Rice_detectMITE', '-cpu',16); runtime=toc; quit).

**GRF** (<https://github.com/bioinfolabmu/GenericRepeatFinder>) is an open-source C++ software on Github. GRF can find multiple types of repeats including terminal inverted repeats (TIRs) and MITEs. GRF requires the CD-HIT software [[5]](https://paperpile.com/c/Ovk0Hl/0a2b) for clustering of repeat candidates. Users can specify the length of TIR/TSD and also the minimum and/or the maximum length of the candidates. We used parameters “-c 0 -t 16 -p 20 --min_space 10 --max_space 10000 --max_indel 0 --min_tr 10 --min_spacer_len 10 --max_spacer_len 10000” for inverted repeats detection, and parameters “-c 1 -t 16 -p 20 --min_space 10 --max_space 10000 --max_indel 0 --min_tr 10 --min_spacer_len 10 --max_spacer_len 10000 --min_tsd 2 --max_tsd 10” for MITE detection. We obtained an unrealistically large number of candidates from both processes, which included 630 Gb and 47 Gb of raw candidates for *grf-main* and *grf-mite*, respectively. We also used default parameters for *grf-mite* to obtain 1.5 Mb of raw candidates. All these results were processed and presented in the benchmarking result (Supplementary Table S1) as GRF-TIR_edu, GRF-mite_edu, and GRF-mite_dft, respectively.

One of the main reasons for the large size of GRF-TIR_edu and GRF-mite_edu raw results is that the majority of these candidates are overlapping and nested within each other, and GRF failed to filter these low-quality candidates. We employed extensive filters to remove overlapping candidates based on the following rules: 1, the minimum length of candidates and their inverted repeat is set to 80 bp and 25 bp (combination of head and tail), respectively. 2, for GRF-mite_edu candidates, target site duplications (TSDs) are required to be present. 3, for candidates with shared start coordinates, retain the candidate with less SNPs and Indels in the inverted repeat region. If multiple start-shared candidates have the same number of mutations in their TIRs, the shortest candidate is retained. 4, for candidates with shared stop coordinates, similar filters were applied as described in 3. 5, for candidates with start coordinates that differ in +/- 1 bp, the candidate with the longer TSD is retained. These filtering steps were implemented in the script “clean_GRF_TIR.pl” in our EDTA toolkit. We also used the script “cleanup_tandem.pl” in our EDTA toolkit to remove tandem repeats in these candidates. After these filters, the size of candidates dropped down to 1.2 Gb and 1.6 Gb for GRF-TIR_edu and GRF-mite_edu, respectively. We further used the script “cleanup_nestedIN.pl” in our EDTA toolkit to remove redundant sequences and nested insertions in the remaining candidates. This step was iterated five times for thorough reduction of redundancy. For MITE candidates, based on a somewhat arbitrary definition, their length should not exceed 600 bp [[26]](https://paperpile.com/c/Ovk0Hl/tWbe), we thus removed candidates that are longer than 600 bp. Finally, the cleaned and non-redundant test library for TIR and MITE are 304 Mb and 10 Mb, respectively.

**miteFinderII** [[27]](https://paperpile.com/c/Ovk0Hl/1Kjh) is an open source C++ software on Github. miteFinderII does not have as many parameter options as others, we used the default parameters (-threshold 0.5). miteFinderII has limited instructions in the README file and refers most questions to the publication.

**MITE-Tracker** [[28]](https://paperpile.com/c/Ovk0Hl/rf9h) is a Python3 package, which needs vsearch [[29]](https://paperpile.com/c/Ovk0Hl/eNIp) for the clustering process. MITE-tracker parameters include the minimum and the maximum length of MITE elements, as well as the minimum and the maximum length of TSD. We used parameters “-w 16 --tsd_min_len 2 --tsd_max_len 9 --mite_min_len 50 --mite_max_len 1000 --task candidates -j Rice” to obtain MITE candidates. Raw candidates contained many false positive sequences with the terminal inverted repeat structure, which may not be useful for TE annotation. The number of clustered MITE candidates is comparable to other programs.

**MUSTv2** [[30]](https://paperpile.com/c/Ovk0Hl/OUyw) is a Perl program that requires BLAST, BLAT [[31]](https://paperpile.com/c/Ovk0Hl/Wkpl), and several Perl packages such as Bioperl [[32]](https://paperpile.com/c/Ovk0Hl/M20w). We used default parameters. The output of MUSTv2 is a .txt file with MITE information such as ID, cluster, coordinates, strand, length, TIR, TSD, and scores. Users need to further process the output .txt file to get the FASTA file of candidate MITEs.

**TIR-Learner** [[33]](https://paperpile.com/c/Ovk0Hl/WdzE) is a Python3 program that uses machine-learning algorithms to facilitate identification and classification of TIR elements. TIR-Learner was originally designed for maize TIR element annotations. In this study, we modified it for the rice genome by using the curated library to train the machine-learning classifier. TIR-Learner integrates the homology-based method and *de novo* machine learning-based method. TIR-Learner uses IRF [[23]](https://paperpile.com/c/Ovk0Hl/5STs) as the TIR structure search engine. We set the minimum TIR=10 and used different length and sequence motifs for identification of TSDs of different TIR superfamilies according to the literature [[34]](https://paperpile.com/c/Ovk0Hl/hMnn). The outputs of TIR-Learner include a gff3 file and a FASTA file. Each entry includes information such as coordinates, superfamily, TIR/TSD sequences/identity, and TE length. To remove potential contaminants from other types of TEs such as LTR retrotransposons, we used non-TIR sequences in the curated library (v6.9.5) to mask TIR-Learner candidates and retain candidates that are longer than 80 bp after removal of masked regions. The test library made of clean candidates is labeled “TIR-Learner_rmLTR” in this study.

We also tried MITE Digger [[35]](https://paperpile.com/c/Ovk0Hl/uLLn), Repetitive Sequence with Precise Boundaries (RSPB) [[36]](https://paperpile.com/c/Ovk0Hl/6eSw), and iMITEdb [[37]](https://paperpile.com/c/Ovk0Hl/UPgv). MITE Digger is a non-open-source Perl program wrapped as a Windows executable. We encountered Perl/Tk errors when running MITE Digger. RSPB has many steps to obtain MITE candidates, with steps 1-3 being automatic and steps 4-6 requiring extensive manual input. The iMITEdb website (<http://gene.cqu.edu.cn/iMITEdb/>) is not accessible.

*Helitron*

**HelitronScanner** [[38]](https://paperpile.com/c/Ovk0Hl/8L8l) is a Java program that utilizes the local combinational variable (LCV) algorithm to identify sequence patterns that are associated with *Helitron* transposons. The program runs on both positive and negative strands of the input genome, and produces candidates with prediction scores that can help to determine the confidence of the prediction. We followed the guidance provided in the maize B73 v4 annotation [[39]](https://paperpile.com/c/Ovk0Hl/ZKvi) and ran HelitronScanner on entire chromosomes without splitting into chunks (-buffer_size 0). We filtered candidates that are not inserted into AT or TT target sites as suggested by [[40]](https://paperpile.com/c/Ovk0Hl/Bkk2) using the script “format_helitronscanner_out.pl” developed in our EDTA toolkit.

Since *Helitrons* tend to capture sequences during their transposition, non-*Helitron* TE sequences and protein-coding sequences could be present. To remove protein-coding sequences, we used the LINE and TIR element transposase database and the MAKER-P plant protein database [[41]](https://paperpile.com/c/Ovk0Hl/rWDd) included in the LTR_retriever package [[11]](https://paperpile.com/c/Ovk0Hl/CT05) to identify non-*Helitron* coding sequences with blastx. Alignment hits with more than 30 aa were retained as real, and if alignments covered ≥ 70% of a *Helitron* candidate, the entire sequence would be discarded. Removal of protein-coding sequence contamination was done using the script “cleanup_proteins” in our EDTA toolkit. To remove non-*Helitron* TE sequences, we used the non-*Helitron* portion of the curated library to mask *Helitron* candidates. The masked sequences were removed and if the remaining sequence was shorter than 100 bp, then the entire sequence was discarded. If the beginning or ending of candidate sequences were masked, then the entire sequence was also discarded due to uncertainty in recognizing the 5'-TC...CTRR-3' *Helitron* structure. Removal of non-*Helitron* TE contamination was accomplished with the script “cleanup_tandem.pl” in our EDTA toolkit with parameters “-misschar N -nc 50000 -nr 0.9 -minlen 100 -minscore 3000 -trf 1 -cleanN 1 -cleanT 1”. Sequence logos of 10 bp flanking and 30 bp terminal sequences were generated using WebLogo 3 (<http://weblogo.threeplusone.com/>) [[42]](https://paperpile.com/c/Ovk0Hl/yPZn).

We also tried HelitronFinder [[43]](https://paperpile.com/c/Ovk0Hl/XGmY), HelSearch [[44]](https://paperpile.com/c/Ovk0Hl/URMT), and a method previously described by Dong *et al.* (2011) [[45]](https://paperpile.com/c/Ovk0Hl/w73a). HelitronFinder seems to be the predecessor of HelitronScanner, however, we could not find the package. HelSearch is a Perl program that relies on blastall which was extremely slow. We split the rice genome into 12 portions (1 file per chromosome) and ran HelSearch in parallel. After three weeks, none of these jobs were finished and we did not find any *Helitron* candidates. Dong *et al.* (2011) developed Perl scripts for identification of *Helitrons*, however, they are not fully automated and not available for public download.

#### Verification of TIR and MITE candidates

TE candidates obtained from detectMITE, GRF-mite_dft, MITE-Hunter, MITE-Tracker, and TIR-Learner were BLAST against the curated library (v6.9.5) to identify any new candidates not contained in the curated library. New candidates were defined as those not confidently matching any sequences in the curated library. Depending on the length of TIRs, short TIRs (i.e., ~10 bp) in these new TIR candidates may still be present in TIR elements in the curated library. We extracted the TIR sequences from new TE candidates based on software output information. For example, in detectMITE, the sequence names are formatted as “>1|6030873|6031020|12|2”, where “12” is the length of the TIR; in GRF, the sequence names are formatted as “>Chr4:7098930:7099636:1m1M10m:TA:3…”, where numbers in “1m1M10m” add up to the length of the TIR (=12). TIR lengths in MITE-Hunter candidates do not have this defined nomenclature to designate the length of the TIR, therefore, 10 bp were extracted from candidate sequences as the minimum length of TIR. TIR sequences were then clustered using the *cd-hit-est-2d* command from CD-HIT [[5]](https://paperpile.com/c/Ovk0Hl/0a2b) with 80% identity threshold. The number of new TIR candidates clustered with the curated library were recorded for each program. For the TIR candidates that possessed novel TIRs, we identified TIR transposon related conserved domains using the NCBI conserved domain database (<https://www.ncbi.nlm.nih.gov/Structure/cdd/>) with default tbastn parameters [[46]](https://paperpile.com/c/Ovk0Hl/rFMe). Redundancy and nesting in novel TIR candidates was removed using the script “cleanup_nestedIN.pl” in our EDTA toolkit, and these non-redundant sequences were used to annotate the rice genome. Copy number of novel TIR candidates in the rice genome were counted by utilizing the script “count_repeats.pl” in our EDTA toolkit. For genomic sequences that cover more than 80% of an TIR candidate, the region was counted as a complete copy, but was otherwise regarded as fragmented.
